# Quinoa produces an insect molting hormone through a plant biosynthetic gene cluster

**DOI:** 10.64898/2026.08.10.743575

**Authors:** Sarah Lam, Anadaisy Aguirre, Marco De La Torre, Asaph Aharoni, Adam Jozwiak

## Abstract

The steroid hormone 20-hydroxyecdysone (20E) controls molting and metamorphosis in arthropods, yet it also accumulates in plants, where its biosynthesis has remained unknown. We show that quinoa (*Chenopodium quinoa*) produces 20E through a seven-gene biosynthetic cluster encoding five cytochrome P450s, a fatty acid hydroxylase-like enzyme (FAH), and a 2-oxoglutarate–dependent dioxygenase (2OGD). Distinct mutant alleles, complementation crosses, and chemical rescue established *FAH* as essential for pathway entry, whereas natural accessions carrying a large cluster deletion lacked 20E and all detectable intermediates. Heterologous reconstruction in *Nicotiana benthamiana* demonstrated that the cluster is sufficient to convert lathosterol to 20E through an unusual FAH-catalyzed C6 oxidation and 2OGD-mediated C5 epimerization. Plants and arthropods therefore evolved distinct enzymatic routes to the same molecule. This pathway provides a blueprint for engineering 20E and defining its broader ecological functions.

## Introduction

Arthropods, which comprise over three-quarters of described animal species, produce and utilize 20-hydroxyecdysone (20E) as their principal molting hormone (*1-3*). Through the EcR/USP receptor, 20E pulses initiate programs directing cuticle synthesis and shedding, metamorphic tissue remodeling, reproduction, and diapause (*3-5*). Because ecdysteroid signaling is essential for arthropod growth, development, and reproduction, it is also a vulnerability that can be used for insect control. Ecdysteroid signaling is exploited by diacylhydrazine insecticides, which activate the ecdysone receptor and induce incomplete and lethal molts (*6,7*). Despite this history, the complete route to 20E remains unresolved in any organism. In the fruit fly (*Drosophila melanogaster*), forward genetics identified the ecdysteroidogenic Halloween enzymes, but some oxidative steps connecting early intermediates remain within the “Black Box” (*8-11*).

Ecdysteroids are also produced by plants. Approximately 5% of surveyed species accumulate phytoecdysteroids, and while more than 500 distinct structures have been described, and 20E is by far the most abundant (*12,13*). Their uneven taxonomic distribution is consistent with repeated gain, loss, or lineage-specific elaboration of phytoecdysteroid metabolism across plant kingdom. The Amaranthaceae are of particular interest in this regard, as several members of the family, including spinach (*Spinacia oleracea*) and quinoa (*Chenopodium quinoa*), accumulate substantial quantities of 20E (*14-17*) (Fig. 1A). Phytoecdysteroids have long been proposed to function as defensive metabolites, disrupting development, feeding, and reproduction of phytophagous insects (*18-21*). Yet plants possess neither an ecdysone receptor nor homologs of the insect *Halloween* genes. Plant 20E biosynthesis therefore cannot represent a simple appropriation of the arthropod pathway; two lineages separated by more than a billion years of evolution have converged upon an identical bioactive molecule by unrelated biochemical routes. Isotope-labeling and metabolite studies suggested precursors and intermediates (*14,22,23*), but no plant 20E pathway had been identified (*24,25*). Thus, the responsible enzyme families, pathway organization, evolutionary dynamics, ecological functions, and potential for heterologous reconstruction remained unresolved.

**Fig. 1.**
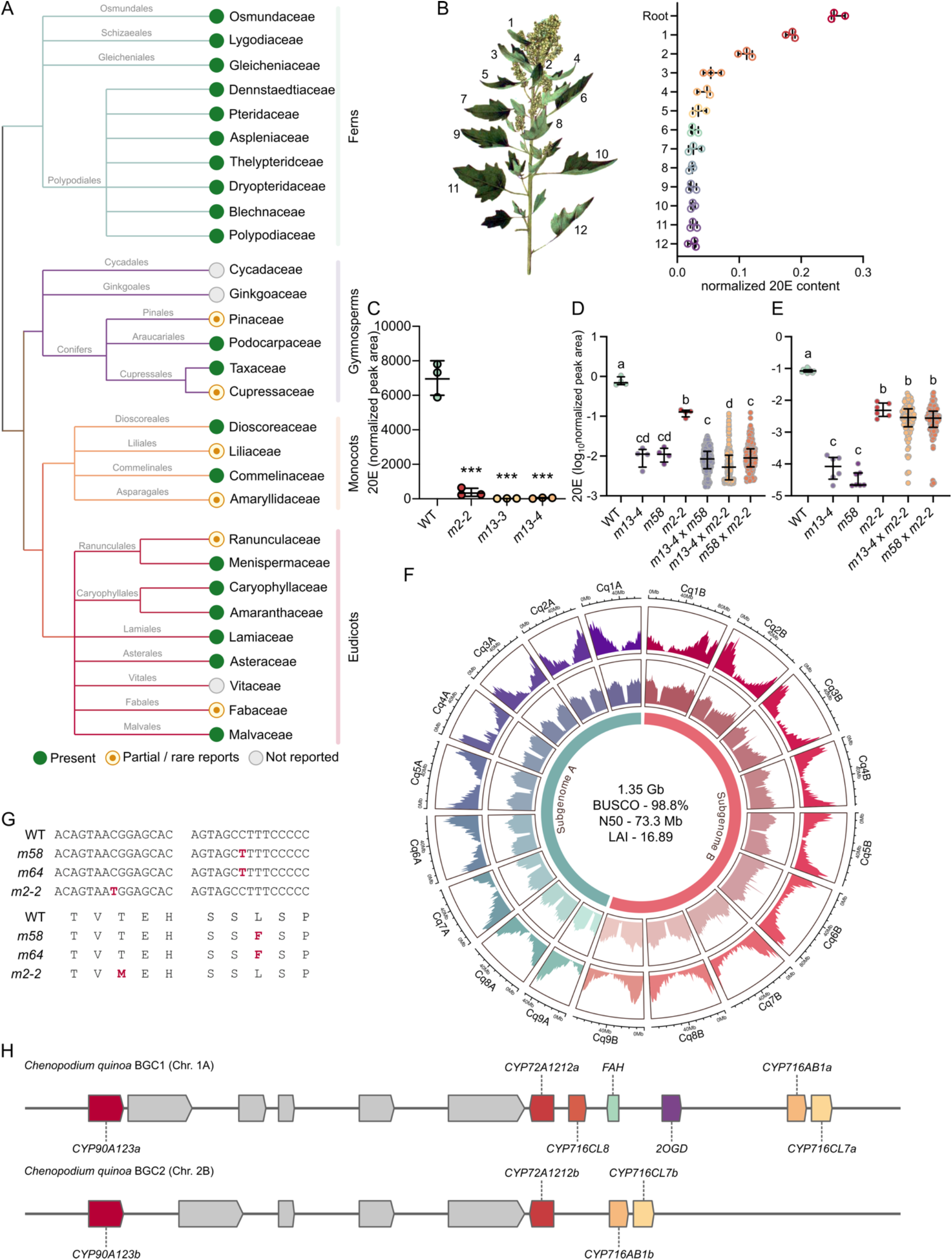
Forward genetics identifies a subgenome-specific 20E biosynthetic cluster in quinoa. (**A**) Simplified phylogeny of representative fern, gymnosperm, monocot, and eudicot families showing the reported occurrence of 20-hydroxyecdysone (20E). Green, present; orange outline, partial or rare reports; gray, not reported. (**B**) Representative quinoa plant showing the sampled root and leaves 1 to 12, ordered from youngest to oldest, and their corresponding normalized 20E abundance (*n* = 3 biological replicates). (**C**) Relative 20E accumulation in wild-type (WT) quinoa and the EMS-derived mutants *m2-2*, *m13-3*, and *m13-4*. (**D**) 20E accumulation in the parental lines and F1 progeny of the indicated complementation crosses. (**E**) 20E accumulation in F2 progeny derived from F1 plants displaying *m2-2-like* 20E levels. In (**B**) to (**E**), points represent individual samples or plants, and horizontal bars show the mean ± SD. In (**C**), \*\**P* < 0.001 relative to WT. In (**D**) and (**E**), groups not sharing a letter differ significantly by one-way ANOVA followed by Tukey’s multiple-comparison test. (**F**) Circos representation of the 18 chromosomes in the de novo RedHead genome assembly. Concentric tracks show gene and repeat density, and assembly statistics are indicated at the center. (**G**) Nucleotide and predicted amino acid changes in *FAH* identified in the EMS mutants: *m58* and *m64* carry the same L168F substitution, whereas *m2-2* carries the distinct T100M substitution; altered nucleotides and residues are highlighted in red. (**H**) Organization of the complete A-subgenome BGC1 on chromosome 1A and the truncated B-subgenome BGC2 on chromosome 2B. Colored arrows indicate 20E biosynthetic genes, gray arrows indicate neighboring genes, and arrow direction denotes transcriptional orientation.

Here, we combine EMS mutagenesis, LC-MS metabolomics, and whole-genome sequencing to identify lesions in a *Fatty-Acid-Hydroxylase-like* (*FAH*) gene required for 20E accumulation. This gene resides in a seven-gene biosynthetic cluster encoding five cytochrome P450s, FAH, and a 2-oxoglutarate-dependent dioxygenase (2OGD). Reconstitution in *Nicotiana benthamiana* showed that the cluster converts lathosterol to 20E through unusual FAH-mediated C6 oxidation and 2OGD-mediated C5 epimerization. Genetic crosses, substrate rescue, and natural accessions lacking the cluster establish its necessity in planta. Comparative genomics reveals assembly of this pathway within an ancient chromosomal neighborhood, followed by subgenome-specific erosion and natural deletions. These findings define the first complete plant 20E pathway and reveal how genomic clustering and catalytic innovation allowed a plant lineage to independently produce an animal steroid hormone.

## Results

### Forward genetics identifies quinoa mutants altered in 20E accumulation

To uncover the genetic basis of plant 20-hydroxyecdysone (20E) biosynthesis, we turned to quinoa (*Chenopodium quinoa*), an unusually powerful system that combines abundant phytoecdysteroid production with agronomic importance and experimental tractability (*15,17,26*). Quinoa is a nutritious and climate-resilient crop that accumulates high levels of 20E, and its monoecious reproductive biology facilitates self-pollination and the recovery of stable mutant lines (*27,28,29*). To determine where 20E accumulates within the plant, we performed LC-MS-based relative quantification across roots and twelve stages of leaf development in 8-week-old plants. Although 20E was detected broadly, its abundance was tissue- and development-dependent, with the highest levels observed in roots and young leaves, and a substantial decline across progressively older leaves (Fig. 1B). This distribution suggests that 20E biosynthesis and accumulation may be spatially separated through long-distance transport, making tissue-specific expression alone insufficient to resolve the pathway unambiguously. We therefore used an unbiased forward-genetic screen to identify mutations affecting 20E production. Approximately 3,600 quinoa seeds were treated with EMS, yielding more than 2,000 M1 plants from which seeds were collected as independent families. We randomly selected 100 M2 families and screened 12 individuals per family by LC-MS for altered 20E accumulation, totaling approximately 1,200 M2 plants (fig. S1,S2). This screen identified two mutant families, family 2 and family 13, bearing plants with strongly reduced or undetectable 20E levels (noise levels) (Fig. 1C, fig. S3-S5). These candidate mutants were advanced by selfing for an additional five to six generations to stabilize the phenotypes. Two mutant lines from family 13, designated *m13-3* and *m13-4*, consistently lacked detectable 20E and were backcrossed to the wild-type Redhead (parental) accession (fig. S6,S7). In contrast, *m2-2* reproducibly accumulated approximately 5-10% of wild-type 20E levels and was maintained as a partial-loss mutant without backcrossing (Fig. 1C). Backcrossed *m13*-derived lines were advanced for three additional generations to yield BC_1_S_3_-58 (*m58*) and BC_1_S_3_-64 (*m64*). Together with the wild-type Redhead, the mutants were selected for whole-genome sequencing to identify mutations responsible for the loss of 20E production.

*Chenopodium quinoa* Willd. (2*n* = 4*x* = 36) is an allotetraploid species containing two subgenomes, A and B, raising the question of whether 20E biosynthesis is redundantly encoded by both subgenomes or primarily controlled by one functional locus. The recovery of strong 20E-deficient mutants from an EMS population, as early as the M2 generation, would be unlikely if complete, redundant pathways were encoded by both subgenomes, suggesting that 20E production may depend predominantly on a single functional subgenomic pathway. To test whether the null and reduced-20E mutations affected the same gene, we performed genetic complementation crosses between *m13-4* and *m2-2*, and between the backcrossed line *m58* and *m2-2* (Fig. 1D,E). If the mutations were in different genes, restoration of wild-type-like 20E levels would be expected in at least some progeny. If they affected the same gene, no full complementation would be expected. In these crosses, seeds were collected from *m13-4* or *m58* plants pollinated by *m2-2*, and more than 100 F1 plants were screened by LC-MS. Most F1 progeny accumulated no detectable 20E, although a small number of individuals accumulated 20E at levels similar to *m2-2*, suggesting residual activity from the *m2-2* allele rather than restoration of wild-type pathway function (Fig. 1D). As a control, a cross between *m13-4* and *m58*, both derived from the same null background, produced no detectable 20E (Fig. 1D). To confirm that the low 20E levels observed in some F1 plants were genetically meaningful, we selfed selected F1 plants and analyzed the F2 progeny. In the F2 generation, approximately 75% of plants accumulated *m2-2*-like low levels of 20E, whereas approximately 25% lacked detectable 20E (Fig. 1E, table S1). This segregation pattern is consistent with mutations occurring within the same gene, with *m2-2* behaving as a hypomorphic allele and *m13-4/m58* behaving as null allele. Together, these results indicate that the null and strongly reduced 20E phenotypes recovered in the EMS screen resulted from recessive mutations at a single genetic locus.

### De novo genome sequencing identifies a candidate lesion in an *FAH-like* gene

Having established that the analyzed 20E-deficient mutants carried recessive mutations affecting the same genetic locus, we next sought to identify the underlying genetic changes. At the time this work was initiated, the only quinoa reference genome was derived from QQ74, a coastal accession, while our EMS population was generated in the RedHead background, a northern highland accession (*27*). Quinoa’s complex domestication history and substantial diversity among coastal and highland accessions could limit analyses based solely on the available reference genome (*27*). Therefore, to aid in mutation discovery, we used PacBio long-read sequencing to generate a highly contiguous genome assembly for the RedHead parental accession and mapped the mutant reads to our new reference assembly. The sequenced mutants included two backcrossed and selfed lines, *m58* (BC_1_S_3_-58) and *m64* (BC_1_S_3_-64), both of which lacked detectable 20E. The wild-type RedHead assembly comprised only 24 contigs, approaching chromosome-level contiguity for the 18 chromosomes of allotetraploid quinoa, with a contig N50 of 73.3 Mb, a total assembly size of 1.35 Gb, BUSCO completeness of 98.8%, and LAI score of 16.89 (Fig. 1F). To prioritize variants most likely to affect 20E biosynthesis, we focused on coding-region mutations predicted to alter amino acid residues. We further highlighted genes with plausible roles in steroid or specialized-metabolite modification, including cytochrome P450s, dioxygenases, reductases, desaturases, oxygenases, hydrolases, and fatty-acid-hydroxylase-like enzymes. This filtering identified 84 biosynthetic genes carrying EMS-type mutations, but one candidate stood out: a *Fatty Acid Hydroxylase-like* gene, which we designated *CqFAH* (hereafter, *FAH*), was the only gene carrying a homozygous nonsynonymous mutation shared by both mutants (Data S1). The two 20E-null lines, which originated from the same M1 family, carried the same C-to-T transition in *FAH*, resulting in a leucine-to-phenylalanine substitution at position 168 (L168F) (Fig. 1G). Subsequent sequencing of *FAH* containing genomic region in *m2-2* identified a distinct EMS-type C-to-T transition that caused a threonine-to-methionine substitution at position 100 (T100M) (Fig. 1G, fig. S8). Together with the expected requirement for oxidative modification of a sterol precursor during 20E biosynthesis, this made *FAH* the leading candidate gene underlying the observed loss of 20E production.

### *FAH* resides in a subgenome-specific biosynthetic gene cluster

Identification of the mutations in *FAH* prompted us to revisit its genomic neighborhood using our newly generated genome assembly of the RedHead variety. We therefore analyzed the genome using plantiSMASH (*30*), which predicted that *FAH* resides within a biosynthetic gene cluster on chromosome 1A (BGC1). The cluster contains seven candidate biosynthetic genes: five cytochrome P450s: *CqCYP90A123a*, *CqCYP72A1212a*, *CqCYP716CL8*, *CqCYP716AB1a*, and *CqCYP716CL7a*, together with *FAH* and a *2-Oxoglutarate-Dependent Dioxygenase* (*Cq2OGD*) (Fig. 1H). The composition of this locus was consistent with the predicted chemistry of 20E biosynthesis, which requires multiple hydroxylation reactions, oxidation at C6, and stereochemical remodeling of the steroid backbone.

Because quinoa is an allotetraploid with A and B subgenomes, we then searched for a corresponding homoeologous cluster in the B subgenome. On chromosome 2B, we identified a related but truncated cluster region (BGC2) containing four cytochrome P450 genes (Fig. 1H). However, this region lacked three genes present in the BGC1, including *FAH*, the *2OGD*, and *CYP716CL8*. Thus, the B-subgenome locus appears to represent an eroded homoeologous cluster rather than a complete redundant pathway. The absence of *FAH* from the BGC2 indicates that the two homoeologous regions are not functionally equivalent: the intact chromosome 1A cluster contains the only enzyme capable of catalyzing the committed early step in 20E biosynthesis, *FAH*.

Thus, despite quinoa’s polyploidy, no B-subgenome *FAH* homolog is available to compensate for loss of the A-subgenome copy. This lack of genetic redundancy provides a mechanistic explanation for why a single EMS-induced lesion in *FAH* was sufficient to cause a severe or complete loss of 20E accumulation. It is also consistent with the observed segregation pattern, which is expected for a phenotype controlled by a single functional locus rather than by two interchangeable homoeologous genes distributed across the two subgenomes.

### Leaf-expressed biosynthetic genes drive root accumulation of 20E

To determine whether the BGC1 could account for 20E production in planta, we next examined transcriptome data across quinoa tissues. The seven BGC1 genes were expressed primarily in leaves and petioles, with little or no expression detected in stems, roots, or floral organs (Fig. 2A; Data S2). This pattern was particularly pronounced for the single-copy genes whose transcript levels were, on average, 50-fold higher in shoots than in roots (Data S2). Genes retained in BGC2 showed a broadly similar spatial expression pattern, with expression mainly in leaves and little or no expression in roots (Fig. 2A). However, their overall transcript abundance was lower than that of the corresponding BGC1 genes. This raised the possibility that BGC2 might retain partial contribution to 20E biosynthesis, but its incomplete gene content and lower expression suggests that it is unlikely to represent the primary functional pathway.

**Fig. 2.**
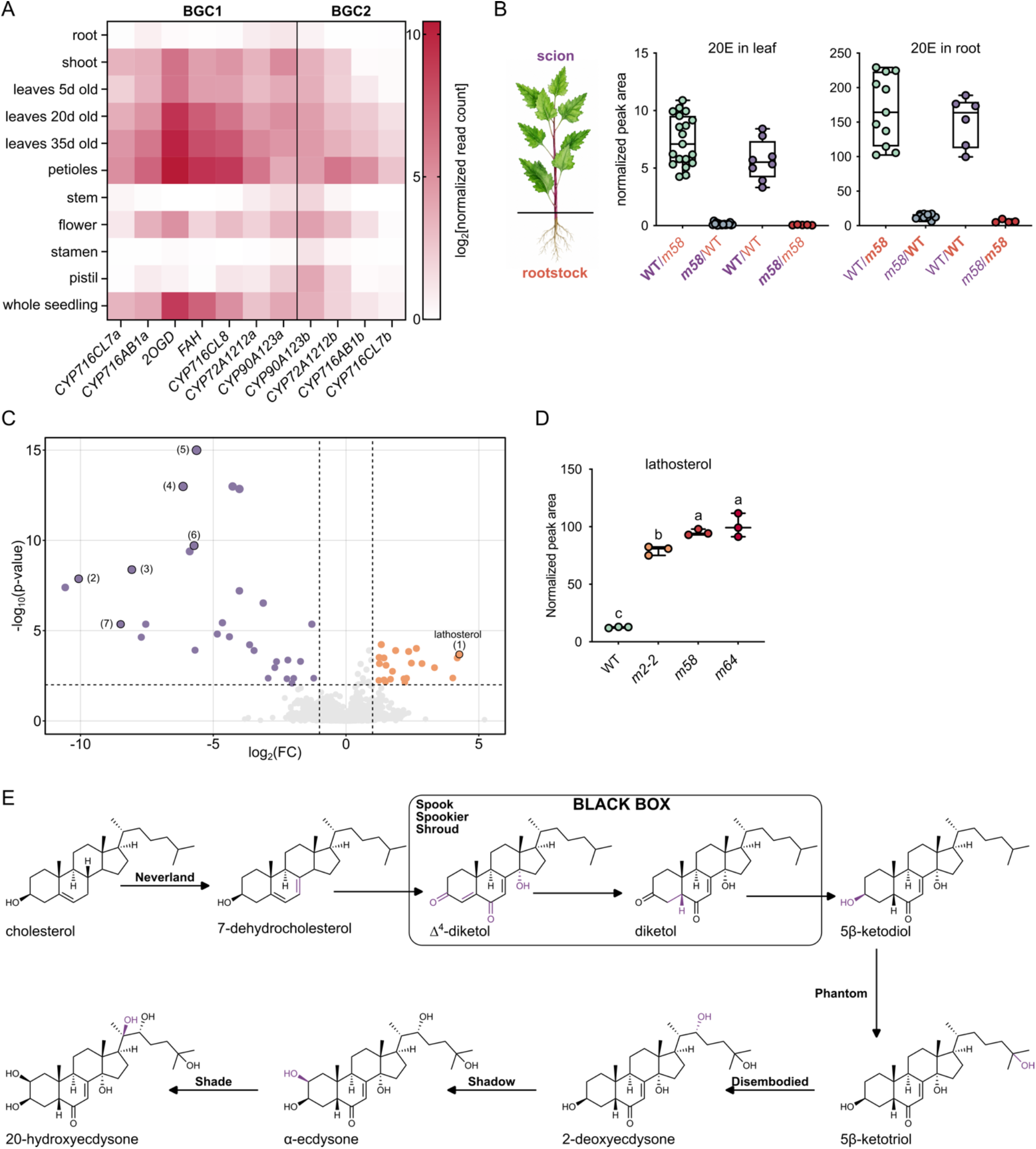
Shoot-localized biosynthesis drives root accumulation of 20E in quinoa. (**A**) Expression of BGC1 and BGC2 genes across quinoa tissues. Values are shown as log_2_-transformed normalized read counts; the vertical line separates genes in the complete BGC1 from those retained in BGC2. (**B**) Reciprocal grafting between wild-type (WT) and 20E-null *m58* plants. Scion genotypes are indicated in purple and rootstock genotypes in orange. 20E abundance was measured separately in leaves and roots; points represent individual grafted plants, and box plots summarize the distributions. (**C**) Differential metabolite accumulation in young leaves of *m58* relative to WT. The volcano plot shows log_2_ fold change versus –log_10_(*P* value); orange and purple points indicate metabolites increased or decreased in *m58*, respectively, and dashed lines denote significance and fold-change thresholds. Lathosterol is highlighted. (**D**) Relative lathosterol abundance in WT and the indicated EMS-derived mutant lines. Points represent biological replicates, and bars show the mean ± SD. Groups not sharing a letter differ significantly by one-way ANOVA followed by Tukey’s multiple-comparison test. (**E**) Ecdysteroid biosynthesis in *Drosophila melanogaster*. Established enzymes are shown in bold, and the unresolved reactions connecting early intermediates to the 5β-ketodiol are enclosed within the Black Box.

This gene expression pattern was unexpected because 20E accumulates broadly in the plant and is especially abundant in roots (Fig. 1B). The lack of correlation between BGC1 transcript abundance and 20E accumulation suggested two possibilities: either additional, root-expressed genes contribute to 20E biosynthesis, or 20E is synthesized in aerial tissues and transported to distal organs. To distinguish between local biosynthesis and long-distance transport, we performed grafting experiments using wild-type and 20E-deficient mutant plants (Fig. 2B). Across multiple grafting configurations, LC-MS analysis showed that roots attached to mutant shoots accumulated only trace amounts of 20E, whereas mutant roots grafted onto wild-type shoots accumulated significantly more 20E (Fig. 2B). These results demonstrate that 20E is transported from the shoot to the root and explain why root 20E abundance does not correspond to local gene expression of the biosynthetic cluster. Consistent with this model, 20E was also highly abundant in root exudates, indicating that shoot-derived 20E is transported belowground and released into the rhizosphere (fig. S9,S10).

Consistently, metabolomic comparison of wild-type and mutant (*m58*) shoots and roots further supported shoot-localized biosynthesis. Mutant and wild-type roots showed little global metabolic divergence, consistent with roots serving primarily as sink or deployment tissues rather than major sites of 20E production (fig. S11). In contrast, young mutant leaves showed pronounced remodeling of steroid metabolism: 20E pathway intermediates were strongly reduced or undetectable, whereas lathosterol was significantly over-accumulated (6.3-fold difference; adjusted *P* = 9.68 × 10^-5^) (Fig. 2C,D). This accumulation pattern identifies lathosterol as the likely entry substrate for quinoa 20E biosynthesis and places FAH at an early committed step in the pathway. The use of lathosterol is notable because insect ecdysteroid biosynthesis depends on dietary sterols, most prominently cholesterol. On the other hand, quinoa appears to route an endogenous Δ^7^ sterol into 20E production (Fig. 2E). Because lathosterol and related Δ^7^ sterols are characteristic metabolites in Caryophyllales (*31*), these findings suggest that plants in this lineage evolved a biochemical route to 20E that is distinct from the arthropod pathway, using a different sterol precursor and a plant-specific set of clustered enzymes.

### Functional reconstruction defines a seven-gene pathway sufficient for 20E biosynthesis

To test whether the BGC1 and BGC2 are sufficient for 20E biosynthesis, we cloned all genes into p3α binary expression vectors and transiently expressed them in *Nicotiana benthamiana* (*32-34*). *Agrobacterium*-mediated co-expression of all seven BGC1 genes in *N. benthamiana* leaves was sufficient to reconstitute 20E production, establishing BGC1 as a complete biosynthetic gene cluster encoding the entire pathway from an endogenous sterol precursor to 20E (Fig. 3A,B,C). Because *N. benthamiana* naturally produces lathosterol as an intermediate in cholesterol biosynthesis, the reconstructed pathway likely used the host-derived substrate without exogenous supplementation. Nevertheless, supplementation with lathosterol increased 20E accumulation approximately five-fold. This increment provides strong functional evidence that lathosterol is the entry substrate for the quinoa pathway and demonstrates that its endogenous availability limits pathway flux in the heterologous host (Fig. 3A,B). To further confirm that lathosterol, rather than cholesterol, serves as the direct precursor of 20E in quinoa, we infiltrated both wild-type quinoa plants and *N. benthamiana* leaves expressing all seven BGC1 genes with stable isotope-labeled cholesterol (cholesterol-d_6_). LC-MS analysis detected no accumulation of labeled 20E in either system, indicating that cholesterol was not incorporated into 20E and does not serve as a direct pathway precursor under these conditions (fig. S12). To determine the contribution of individual pathway genes to 20E biosynthesis, we systematically omitted each of the seven candidate genes during transient reconstitution in *N. benthamiana*. These assays showed that removal of any single gene abolished detectable 20E accumulation, indicating that all seven genes are required for complete 20E production in this heterologous system (Fig. 3D). The absence of detectable complementation by endogenous *N. benthamiana* enzymes makes this system well suited for unambiguous functional assignment of each pathway component. We next tested the retained genes from the eroded B-subgenome cluster by replacing the corresponding BGC1 cytochrome P450s with their BGC2 counterparts. Consistent with their homoeologous origin, all genes retained in both clusters are highly conserved, sharing 92-97% amino acid sequence identity (fig. S13). These BGC2 enzymes were functional and supported 20E production, but with approximately 40% of the efficiency observed with the BGC1 enzymes under our assay conditions (Fig. 3E). Thus, although the B-subgenome cluster is incomplete, its retained P450 genes remain catalytically active and may contribute partially to 20E metabolism when supplied with the missing pathway components.

**Fig. 3.**
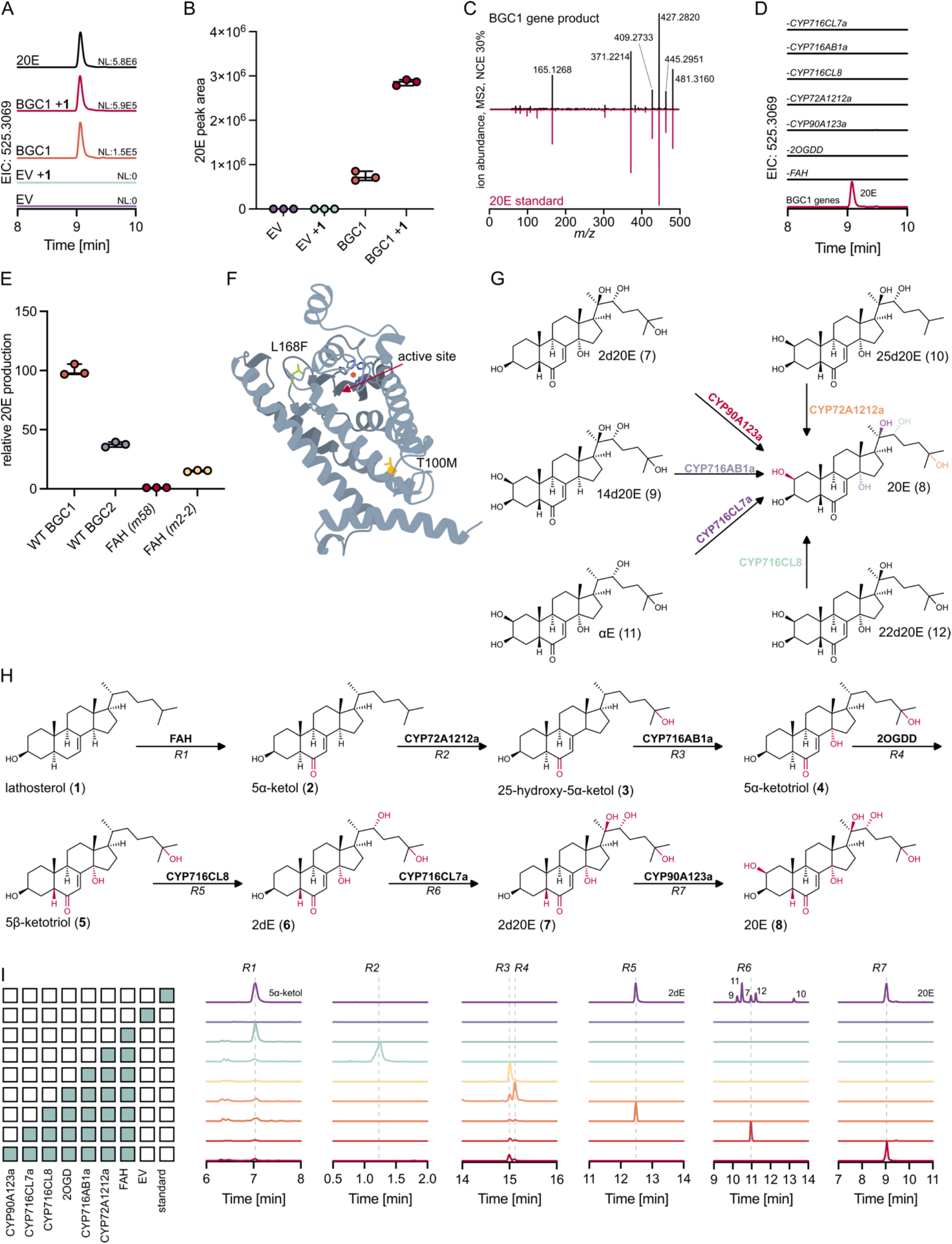
Heterologous reconstruction defines the complete quinoa pathway from lathosterol to 20E. (**A**) Extracted-ion chromatograms (EICs) for the 20E formate adduct, *m/z* 525.3069 [M + formate - H]^-^, in *Nicotiana benthamiana* leaves expressing empty vector (EV) or all seven BGC1 genes, with or without lathosterol supplementation (+1). An authentic 20E standard is shown in black. (**B**) 20E abundance in the samples shown in (A). Points represent biological replicates, and bars show the mean ± SD. (**C**) Mirror plot comparing the MS^2^ spectrum of 20E produced by BGC1 expression with that of an authentic 20E standard at normalized collision energy 30%. (**D**) Gene-omission analysis of BGC1. EICs show that detectable 20E production required coexpression of all seven genes; the omitted gene is indicated above each trace. (**E**) Relative 20E production by the complete BGC1 pathway, a pathway in which P450s retained in BGC2 were substituted for their BGC1 homologs, and pathways containing the *m58* or *m2-2* FAH alleles. Values are normalized to the wild-type BGC1 pathway; points represent biological replicates, and bars show the mean ± SD. (**F**) AlphaFold3 model of FAH showing the predicted active-site region and the positions of the L168F and T100M substitutions identified in *m58*/*m64* and *m2-2*, respectively. (G) Functional assignment of pathway P450s by feeding monodehydroxylated 20E derivatives to *N. benthamiana* expressing individual enzymes. Arrows indicate conversion of each substrate to 20E and identify CYP90A123a, CYP716AB1a, CYP716CL7a, CYP716CL8, and CYP72A1212a as the C2-, C14-, C20-, C22-, and C25-hydroxylases, respectively. (**H**) Complete quinoa 20E biosynthetic pathway from lathosterol (1) to 20E (8). Enzymes are shown above each reaction, *R1* to *R7* denote the sequential reactions analyzed in (I), and newly introduced functional groups are highlighted in red. (**I**) Stepwise and combinatorial reconstruction of the pathway in *N. benthamiana*. The matrix indicates the combinations of expressed genes, with filled squares denoting included constructs. EICs show accumulation of the products assigned to reactions *R1* to *R7*; dashed lines indicate the retention times of the corresponding pathway products or authentic standards.

We then used the *N. benthamiana* heterologous expression system to determine whether the quinoa mutations directly impaired FAH function. Substituting the wild-type enzyme with the *m58* allele abolished 20E production, whereas the *m2-2* allele retained only approximately 10% of wild-type activity (Fig. 3E). These effects closely matched the 20E phenotypes observed in quinoa, where *m58* lines lacked detectable 20E and *m2-2* retained only strongly reduced accumulation (Fig. 1C). Thus, the L168F substitution in *m58* nearly eliminates FAH activity, whereas the T100M substitution in *m2-2* produces a hypomorphic allele with residual activity (Fig. 1C,3E). AlphaFold 3 structural models further supported these functional differences: L168F lies close to the predicted active-site, consistent with a severe effect on catalysis or substrate positioning. On the other hand, T100M is positioned farther from the active site in a predicted transmembrane region, where it may affect substrate access, membrane positioning, or retention of the sterol substrate, affecting 20E biosynthesis to a lesser degree (Fig. 3F).

### Biochemical reconstruction resolves the pathway from lathosterol to 20E

To assign the regiospecific activity of each pathway P450 and 2OGD, we first expressed the candidate enzymes individually in *N. benthamiana* and supplied the corresponding monodehydroxylated 20E analog lacking a single hydroxyl group at C2, C14, C20, C22, or C25.

In each assay, the supplied substrate was converted directly to 20E, establishing CYP90A123a, CYP716AB1a, CYP716CL7a, CYP716CL8, and CYP72A1212a as the C2-, C14-, C20-, C22-, and C25-hydroxylases, respectively (Fig. 3G, fig. S14). By contrast, 2OGD did not convert any of the monodehydroxylated substrates to 20E, indicating that it does not catalyze one of these hydroxylation steps and likely performs a different reaction (fig. S14). Because these feeding assays defined the position modified by each P450 but not the order of the reactions in the native pathway, we next performed stepwise and combinatorial reconstruction from lathosterol in *N. benthamiana*. The pattern of intermediate accumulation and product formation identified the predominant productive sequence under the reconstruction conditions. FAH first converts lathosterol (**1**) to 5α-6-oxolathosterol, also known as 5α-ketol (**2**) (fig. S15). CYP72A1212a then introduces the C25 hydroxyl group to generate 25-hydroxy-5α-ketol (**3**), followed by C14 hydroxylation by CYP716AB1a to produce 5α-ketotriol (**4**) (Fig. 3G,H; fig. S16). The 2OGD subsequently catalyzes C5 epimerization, converting 5α-ketotriol (**4**) to 5β-ketotriol (**5**) (fig. S16,S17). CYP716CL8, CYP716CL7a, and CYP90A123a then act sequentially to introduce hydroxyl groups at C22, C20, and C2, yielding 2-deoxyecdysone (2dE; **6**), 2-deoxy-20E (2d20E; **7**), and finally 20E (**8**), respectively (Fig. 3G,H). Although 20E production was efficient in *N. benthamiana*, several pathway intermediates and 20E itself were glycosylated by endogenous glycosyltransferases, likely reducing overall 20E yield (fig. S18). To determine whether the absence of 20E in quinoa mutants resulted specifically from disruption of the first committed, FAH-catalyzed step, rather than from defects elsewhere in the pathway, we infiltrated young *m58* leaves with 5α-ketol, 5β-ketol, and monodehydroxylated 20E intermediates. Each supplied intermediate was converted to 20E, demonstrating that the downstream pathway remains functional (fig. S19). Analysis of WT quinoa leaves further supported the inferred reaction order. The most abundant monodehydroxylated intermediate was 2d20E (**7**), consistent with C2 hydroxylation being the terminal step, while the presence of 2dE (**6**) supports C20 hydroxylation immediately preceding it (fig. S20). Together, direct substrate-feeding assays, combinatorial reconstruction, chemical complementation of the *m58* mutant, and detection of pathway intermediates in planta establish the enzymatic activities and reaction sequence from lathosterol to 20E. Collectively, these findings define a previously unknown route to 20E that is fundamentally different from the arthropod pathway in both precursor usage and enzyme composition. The recruitment of an endogenous Δ^7^ sterol, an FAH-like C6 oxidase, a 2OGD epimerase, and multiple CYP716 enzymes strongly supports independent evolution of 20E biosynthesis in quinoa and suggests that chemically convergent 20E production in distantly related plant lineages arose through distinct enzymatic solutions.

### Cluster assembly and erosion shaped quinoa 20E biosynthesis

To determine when and how the quinoa 20E cluster arose, and how its architecture changed during subsequent evolution, we compared its genomic organization across Caryophyllales. Using the ∼230-kb, 25-gene BGC1 interval on chromosome 1A of the quinoa genome as an anchor, synteny analysis across 17 genomes revealed conserved collinearity of the flanking non-biosynthetic genes throughout Amaranthaceae, Caryophyllaceae, Droseraceae, and Nepenthaceae (Fig. 4A). This broad conservation demonstrates that the underlying chromosomal neighborhood substantially predates assembly of the 20E pathway and provides a framework for tracing the lineage- and subgenome-specific remodeling of the cluster. Against this ancient syntenic backbone, however, all seven biosynthetic genes were retained as a compact, co-linear cassette only in the sampled Chenopodioideae, including *Chenopodium* species and spinach. By contrast, more distantly related Caryophyllales genomes retained homologs of the constituent enzyme families at dispersed, nonsyntenic positions. Importantly, the absence of a BGC1-like locus does not indicate an absence of ecdysteroid biosynthesis: 20E and multiple related ecdysteroids have been documented in several *Silene* (Caryophyllaceae) and *Amaranthus* (Amaranthaceae) species (*16*), yet the corresponding enzyme homologs in the genomes examined here were not organized into a comparable seven-gene cluster (Fig. 4A). This contrast suggests that related ecdysteroid chemistry can arise from distinct genomic architectures, ranging from the compact Chenopodioideae cluster to more dispersed enzyme complements in other Caryophyllales lineages. These results indicate that the pathway enzymes did not originate *de novo* with the cluster. Rather, they were recruited from a pre-existing genomic repertoire and consolidated within a conserved chromosomal region, likely through gene duplication and relocation. The shared position, gene content, and compact organization of the cassette in spinach and *Chenopodium* support inheritance from a single assembly event in their common ancestor rather than independent formation in each lineage. However, an older cluster followed by complete loss from all sampled outgroups cannot be formally excluded. Subsequent remodeling of this ancestral locus is particularly evident in quinoa. The BGC1 retains the complete seven-gene pathway, whereas BGC2 retains only *CYP90A123b*, *CYP72A1212b*, *CYP716AB1b*, and *CYP716CL7b* and lacks recognizable pathway-associated copies of *FAH*, the *2OGD*, and *CYP716CL8* (Fig. 1H,4B). Examination of related *Chenopodium* genomes further resolved the timing of this degeneration. BGC1-like loci were detected in the diploid relatives *C. pallidicaule* and *C. suecicum*, representing the A- and B-subgenome lineages, respectively, while the closely related allotetraploid *C. berlandieri* appeared to retain intact clusters on both its A- and B-derived subgenomes (*27*) (Fig. 4B, fig. S21). These comparisons indicate that the incomplete structure of quinoa BGC2 is not an ancestral property of the B lineage, but instead reflects subgenome-specific erosion during the evolution of cultivated quinoa or its immediate progenitor.

**Fig. 4.**
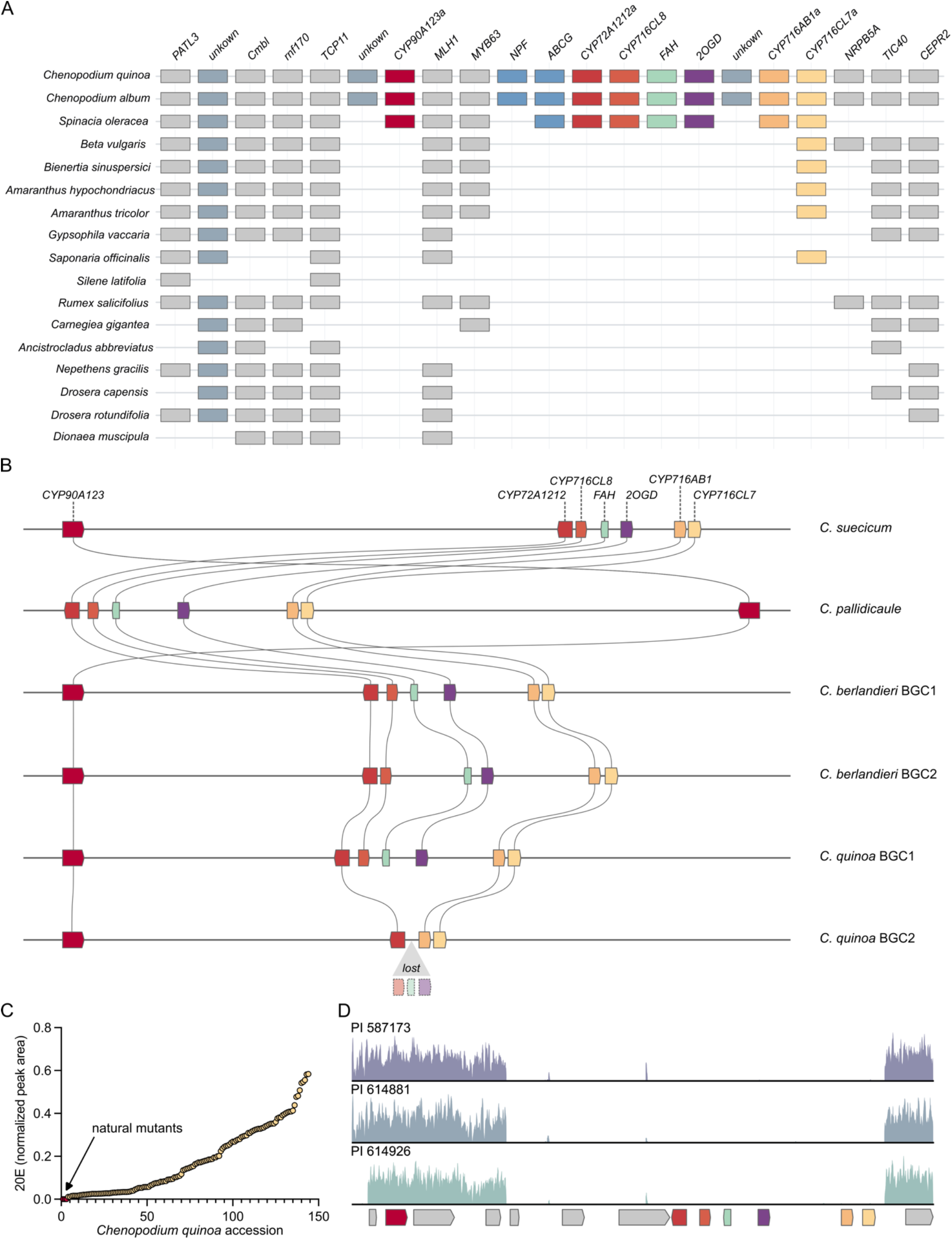
Comparative genomics reveals assembly, erosion, and natural loss of the quinoa 20E cluster. (**A**) Conservation of the BGC1-homologous genomic interval across 17 representative Caryophyllales genomes. Columns correspond to genes in the quinoa chromosome 1A interval, and boxes indicate orthologs retained within the syntenic region. Colored boxes denote the seven 20E biosynthetic genes, using the same colors throughout the figure; gray and blue-gray boxes denote neighboring nonbiosynthetic genes. The complete seven-gene cassette is retained only in the sampled Chenopodioideae, whereas more distant genomes preserve portions of the surrounding syntenic backbone but lack a comparable cluster. (**B**) Synteny of 20E cluster loci in diploid *Chenopodium* relatives, allotetraploid *C. berlandieri*, and quinoa. Curved lines connect homologous pathway genes, and colors identify the corresponding enzymes. Complete clusters are retained in *C. suecicum*, *C. pallidicaule*, and both subgenomes of *C. berlandieri*, whereas quinoa BGC2 lacks *FAH*, *2OGD*, and *CYP716CL8*, shown below the locus as lost genes. (**C**) Normalized 20E abundance across approximately 150 quinoa accessions ordered by increasing accumulation. Each point represents three biological replicates of one accession; the arrow indicates three natural accessions with no detectable 20E. (**D**) Illumina read-depth profiles across BGC1 in the three naturally 20E-null accessions. All three show a shared loss of coverage spanning most of the cluster, while coverage is retained over the flanking regions and the distal *CYP90A123a* locus. The BGC1 gene model is shown below, with pathway genes colored as in (A) and (B).

Natural variation within quinoa accessions provides additional evidence that the intact A-subgenome cluster is the principal functional source of pathway flux. LC-MS screening of approximately 150 quinoa accessions identified three accessions with no detectable 20E: PI614881 and PI587173, both originating from Argentina, and PI614926, originating from Bolivia (Fig. 4C, fig. S22,S23). Because the genetic and biochemical data implicated BGC1 as the active biosynthetic locus, we attempted to amplify and sequence genes across the cluster in these natural 20E-null accessions. Most BGC1 genes failed to amplify, whereas *CYP90A123a*, located at the distal edge of the cluster, remained detectable (fig. S24). Whole-genome resequencing confirmed that this pattern resulted from a large deletion spanning the majority of BGC1 (Fig. 4D). The deletion boundaries appeared identical in all three accessions, consistent with inheritance of the same structural variant from a common ancestral deletion event rather than three independent losses. In contrast, the architecture of the B-subgenome BGC2 locus was indistinguishable from that of 20E-producing accessions and showed no additional degeneration (fig. S25). The 20E-null phenotype in these natural accessions can therefore be attributed specifically to deletion of the A-subgenome BGC1, rather than to additional erosion of BGC2. Consistent with elimination of the functional pathway, the three accessions lacked not only 20E but also detectable biosynthetic intermediates, indicating that the deletion abolishes pathway flux rather than blocking a late tailoring reaction (fig. S26). Together, these results define the quinoa-type 20E cluster as a lineage-restricted consolidation of evolutionarily older biosynthetic components within an ancient syntenic neighborhood, followed by differential retention of homoeologous copies, selective erosion of BGC2, and natural near-complete deletion of the functional BGC1 locus in some accessions. The 20E cluster therefore behaves as a dynamic genomic module whose physical organization promotes coordinated inheritance of the pathway but also permits its rapid loss through a single structural event.

### Stepwise recruitment and catalytic reinvention shaped the 20E cluster

Phylogenetic analyses demonstrated that BGC1 is a mosaic of enzyme genes derived from distinct and substantially older evolutionary lineages. CYP90A123a and its BGC2 homolog CYP90A123b formed a maximally supported cluster-associated lineage shared with spinach and *Chenopodium album* but distinct from a broadly retained CYP90A clade containing homologs from *Beta*, *Bienertia*, *Amaranthus*, spinach, and *Chenopodium* (Fig. 5A). This topology identifies CYP90A123 as a derived paralog and is consistent with duplication of an ancestral Amaranthaceae CYP90A gene, substantial divergence and differential retention of one copy in the sampled Chenopodioideae, and its subsequent recruitment into the ancestral 20E cluster. The long branch leading to the CYP90A123 lineage suggests that much of this sequence specialization preceded the spinach-*Chenopodium* divergence, although the tree alone cannot establish whether C2-hydroxylase activity evolved before or after physical incorporation into the cluster (Fig. 5A).

**Fig. 5.**
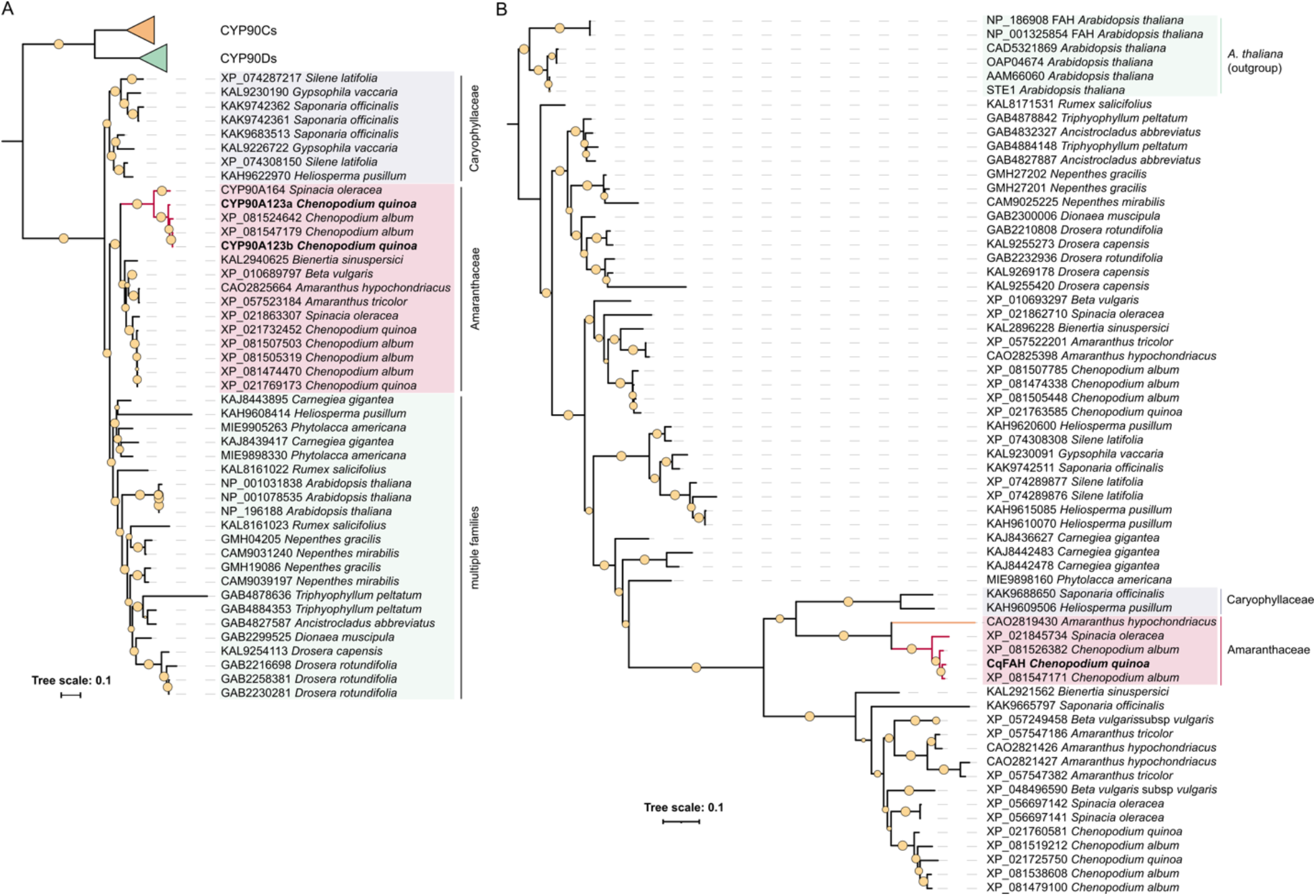
Phylogenetic analyses reveal recruitment of derived CYP90A and FAH paralogs into the 20E cluster. (**A**) Maximum-likelihood phylogeny of CYP90A proteins from representative Caryophyllales species, rooted with CYP90C and CYP90D sequences. Quinoa CYP90A123a and its BGC2 homolog CYP90A123b form a distinct cluster-associated lineage with homologs from spinach and *Chenopodium album*, separate from the broadly retained CYP90A clade. (**B**) Maximum-likelihood phylogeny of FAH-like proteins across Caryophyllales, rooted with *Arabidopsis thaliana* homologs. Quinoa FAH falls within a distinct lineage containing homologs from Amaranthaceae and Caryophyllaceae, consistent with recruitment of a pre-existing paralog rather than recent duplication within the cluster. Functionally characterized quinoa enzymes are shown in bold, shaded regions indicate major taxonomic groups or clades, and node symbols denote supported branches. Scale bars indicate amino acid substitutions per site.

CYP72A1212a likewise belonged to a strongly supported spinach-*Chenopodium* clade nested within a broader CYP72A1212-like lineage that included *Silene* and *Heliosperma* proteins, indicating that the ancestral cluster recruited a pre-existing CYP72A paralog whose origin predates the divergence of Amaranthaceae and Caryophyllaceae (fig. S27). Two quinoa-*C. album* sublineages were retained within this clade, consistent with the presence of homoeologous *CYP72A1212* genes in BGC1 and BGC2 (fig. S28).

The three pathway CYP716 enzymes had still more complex origins. CYP716AB1 belonged to an ancient Caryophyllales lineage distinct from the CYP716CL branch, whereas CYP716CL7 and CYP716CL8 formed strongly supported sister clades containing *Heliosperma*, spinach, and *Chenopodium* homologs (fig. S28). This taxonomic distribution places the CYP716CL7/CL8 duplication before the divergence of Amaranthaceae and Caryophyllaceae and indicates that two already-diverged CYP716CL paralogs were incorporated into the ancestral Chenopodioideae cluster rather than generated through a recent cluster-localized expansion.

FAH similarly belonged to one branch of an ancient Caryophyllales duplication that preceded the Amaranthaceae-Caryophyllaceae split. Its placement within a strongly supported Chenopodioideae cluster-associated clade nested within a broader lineage containing proteins from *Amaranthus hypochondriacus*, *Heliosperma pusillum*, and *Saponaria officinalis* (all producing phytoecdysteroids) demonstrates that the cluster recruited a pre-existing *FAH*-like paralog rather than generating it through a recent local duplication (Fig. 5B).

Finally, the pathway 2OGD was nested within an older Caryophyllales LBO/SRG1/JOX-like radiation but formed a distinct pathway-associated lineage with spinach, *C. album*, and an additional Chenopodioideae-related sequence (fig. S29). The absence of second quinoa proteins from the immediate FAH, 2OGD, and CYP716CL8 lineages provides independent phylogenetic support for loss of their homologs from BGC2. Collectively, these trees show that the complete pathway did not arise through expansion of a single ancestral locus but was assembled stepwise from paralogs drawn from several ancient enzyme families.

Recruitment of these ancient paralogs was accompanied by extensive catalytic reinvention: FAH, a membrane-bound non-heme iron hydroxylase/desaturase-family protein, forms a C6 ketone on lathosterol rather than catalyzing canonical desaturation or hydroxylation; a 2OGD performs redox-neutral C5 epimerization; three CYP716s install C14, C20, and C22 hydroxyls on an ecdysteroid scaffold; and a derived CYP90A completes 20E formation by C2 hydroxylation (Fig. 3H). Together, these activities establish the quinoa cluster as both an assembly of ancient paralogs and a source of previously unrecognized plant enzymatic chemistry.

## Discussion

We define a complete plant route from an endogenous sterol to 20-hydroxyecdysone (20E), resolving a question posed since phytoecdysteroids were first described more than half a century ago (*12,35*). The pathway is not a variant of the arthropod route: quinoa and insects share neither precursor, enzyme families, nor genomic organization. Insects convert dietary cholesterol through the Rieske oxygenase Neverland and an unresolved series of “Black Box” reactions leading to the 5β-ketodiol (*9-11*). Quinoa instead converts lathosterol, an endogenous Δ^7^ sterol abundant in Caryophyllales, using a fatty-acid-hydroxylase-like oxidase, CYP72A, three CYP716 enzymes, a derived CYP90A paralog, and a 2OGD epimerase encoded within a single ∼230-kb cluster. Identical products therefore conceal complete biochemical independence. The selection of lathosterol helps explain how this pathway became accessible: every ecdysteroid contains a Δ^7^-6-one enone, and lathosterol already supplies the Δ^7^ bond, leaving only C6 oxidation to complete this defining chromophore. Insects must first generate the unsaturation from cholesterol through Neverland (*10*), whereas Chenopodioideae simply inherit it from primary sterol metabolism. We therefore propose that abundant Δ^7^ sterols provided a permissive substrate pool that lowered the biochemical threshold for pathway assembly. This predicts that 20E-producing lineages in which Δ^7^ sterols are not prominent, including *Ajuga* and ferns, enter the pathway at a different oxidation state and through different enzymes (see accompanying paper by Park et al.).

The first committed step illustrates how thoroughly the pathway was improvised from existing parts. FAH belongs to the membrane-bound di-iron fatty acid hydroxylase superfamily, whose characterized members desaturate sterols or hydroxylate acyl chains during sphingolipid biosynthesis (*36,37*). Quinoa FAH does neither: it converts lathosterol to 5α-6-oxolathosterol, installing a C6 ketone and completing the ecdysteroid enone in one step. Although plants already use CYP85A enzymes for steroid C6 oxidation in brassinosteroid biosynthesis (*38,39*), the 20E pathway independently recruited a desaturase-family protein for the same positional transformation. The P450s reveal an equally striking reassignment of catalytic roles. CYP716 enzymes are best known for oxidizing β-amyrin and related triterpene scaffolds in saponin biosynthesis (*40,41*), yet three quinoa CYP716s hydroxylate an ecdysteroid nucleus at C14, C20, and C22, activities that, to our knowledge, are unprecedented among characterized plant P450s in this family. CYP90 enzymes normally catalyze C3, C22, and C23 oxidations in brassinosteroid biosynthesis (*42-44*), whereas CqCYP90A123a performs the terminal C2 hydroxylation, a reaction not previously reported for this family.

The most unexpected enzyme is the 2OGD. Every ecdysteroid has a 5β-configured, *cis*-fused A/B ring junction, yet the origin of this stereochemistry remains unresolved in insects and lies within the “Black Box” (*11*). Known pathways typically build the 5β configuration by reducing a Δ^4^ intermediate, as in *Digitalis* cardenolide biosynthesis and mammalian bile acid formation (*45,46*). Quinoa instead directly converts 5α-ketotriol to 5β-ketotriol without a Δ^4^ intermediate or net change in oxidation state. The recently described *Digitalis* 2OGD S14βH provides a plant precedent for controlling steroid stereochemistry, but couples inversion to C14 hydroxylation through substrate reorientation and opposite-face hydroxyl rebound (*47*). The quinoa enzyme instead achieves net epimerization without product hydroxylation, potentially through opposite-face hydrogen return. Bacterial CarC and SnoN provide mechanistic precedents for such Fe/2OG-dependent stereoinversion (*48-50*). To our knowledge, this is the first 2OGD shown to mediate redox-neutral epimerization of an intact steroid nucleus, offering a potential biocatalyst for stereochemical editing of steroid scaffolds.

Our spatial data raise a second question: why synthesize 20E in shoots and export it belowground? The cluster is expressed mainly in young leaves, yet 20E accumulates in roots and surrounding soil. Spinach provides a relevant precedent: wounding and herbivory induce phytoecdysteroid production, 20E moves toward damaged roots (*51*), and these metabolites impair plant-parasitic nematodes by causing abnormal molting, reduced invasion, and death (*20*). Together, these findings support a centralized source with belowground delivery rather than passive leaf storage. Shoot-localized synthesis may exploit greater precursor availability while avoiding duplication of the pathway in roots. However, soil release could reflect either defense against belowground pests or disposal of excess metabolite. The 20E mutants and three natural null accessions now make these alternatives genetically testable through nematode challenge, root herbivory, and rhizosphere profiling, converting a long-standing hypothesis into an experiment.

The cluster’s evolutionary history is as unusual as its chemistry. Its genes are ancient: *FAH*, *CYP716CL7/8*, *CYP72A1212*, and the pathway *2OGD* are derived from Caryophyllales lineages predating the Amaranthaceae-Caryophyllaceae split. Their arrangement is younger, restricted to the sampled Chenopodioideae and shared by spinach and *Chenopodium*. The principal innovation was therefore genomic consolidation of pre-existing paralogs rather than invention of enzyme families. *Silene* and *Amaranthus* species produce ecdysteroids without a comparable cluster (*12*), showing that physical clustering is not required for the chemistry. Its advantage must instead lie in properties such as co-inheritance, coordinated regulation, or containment of intermediates (*52-54*). Like the *Arabidopsis* thalianol locus, the quinoa cluster illustrates how plant biosynthetic gene clusters arise through duplication, relocation, and neofunctionalization of native genes (*54-57*). Unlike the largely fixed thalianol cluster, the 20E pathway occurs in complete, eroded, deleted, and dispersed states across Caryophyllales, exposing both the advantages and fragility of physical linkage. Because the thalianol cluster shapes the *Arabidopsis* root microbiota (*58*) and quinoa exports 20E into the rhizosphere, the selective value of linkage in both systems may ultimately be realized belowground.

The same linkage that favors co-inheritance also enables wholesale loss. Quinoa carries an eroded BGC2, while three natural accessions share a large BGC1 deletion. BGC2 erosion may reflect biased fractionation following hybridization, with one homolog becoming dispensable. Natural BGC1 loss may instead reflect relaxed selection under cultivation, with the metabolic cost of diverting Δ^7^ sterols from membranes or potential antagonistic effects on beneficial soil organisms. Domestication-associated loss of seed saponins in quinoa through *TSARL1* provides a relevant precedent: both cases involve loss of defensive chemistry, although one is regulatory and tissue-specific and the other structural and complete (*27,59*). These hypotheses can be distinguished by tracing the deletion haplotype across Andean germplasm and testing null lines under defined biotic pressures. The restriction of all three null accessions to Argentina and Bolivia further favors inheritance within a shared ecological or agronomic context over recurrent independent deletion.

Quinoa therefore produces an arthropod hormone itself, not merely a structural mimic, through a pathway sharing no enzyme with the animals that use it. Because the compact pathway is sufficient to produce 20E in *N. benthamiana*, it could be transferred to other biological systems. Its unusual enzymes also provide new tools for introducing position-specific oxidation and stereochemical changes into steroid scaffolds. The broader question is now how many times, and through how many enzymatic solutions, have plants independently evolved 20E? The quinoa cluster provides the first complete answer for one lineage and a template for finding the remaining.

## Acknowledgments

We thank Ryan Nett for generously providing standards of 20E pathway intermediates. Seeds of quinoa accessions PI 614881, PI 587173, and PI 614926 were obtained from the USDA National Plant Germplasm System quinoa collection. We would like to thank our undergraduate researchers, John Amaya and Damandeep Kaur, for their contributions towards data collection.

## Author contributions

Conceptualization: AJ, SL, Aah

Methodology: SL, AJ

Investigation: SL, AAg, MDLT, AJ

Visualization: SL, AJ

Funding acquisition: AJ, Aah

Project administration: AJ

Supervision: AJ, Aah

Writing – original draft: AJ

Writing – review & editing: AJ, SL, AAh

## Data, code, and materials availability

All data needed to evaluate the conclusions in the paper are present in the main text, supplementary materials, or public repositories. Raw genome sequencing reads and genome assemblies for wild-type RedHead quinoa and mutant-derived lines will be available at NCBI under accession numbers [XXXX] upon manuscript acceptance. Plasmids, seeds, mutant lines, and other biological materials generated in this study are available from the corresponding author upon reasonable request, subject to availability, phytosanitary regulations, institutional approvals, and material transfer agreements.

## Supplementary Materials

Materials and Methods

Figs. S1 to S29

Tables S1 to S2

References (*60*–*110*)

Data S1 to S3

